# SpliSync: Genomic language model-driven splice site correction of long RNA sequencing reads

**DOI:** 10.64898/2026.07.04.736518

**Authors:** Wui Wang Lui, Liliana Florea

## Abstract

Long RNA sequencing reads are rapidly replacing short reads in transcriptomic analyses, enabling full-length transcript sequencing and better identification of isoforms, alternative splicing events, and other transcript variants. However, their higher sequencing error rates can cause misalignments, especially at splice junctions, reducing the accuracy of transcript reconstruction and analysis. We developed SpliSync, a genomic language model-driven method for splice site correction that integrates a pre-trained genomic sequence model (HyenaDNA), alignment data, and a U-net architecture to predict splice sites at nucleotide resolution. SpliSync substantially improved the precision of RNA long-read alignments by 27%-194% across diverse datasets and consistently outperformed competing tools. As a preprocessing step, it increased alternative splicing detection accuracy by 26%-330%. In contrast, its benefit for transcript reconstruction was limited, likely due to the tools’ built-in correction mechanisms. The code was developed in Python using the PyTorch package, and is freely available at https://github.com/splicebox/SpliSync.

## INTRODUCTION

Long-read RNA sequencing has revolutionized transcriptomic research, enabling the sequencing of full-length transcripts and better identification of isoforms, alternative splicing events, and other transcript variants (1). Technologies such as PacBio and Oxford Nanopore can produce enough reads to enable surveying the transcriptome. However, the sequence quality is below that of RNA-seq sequences, with 3-10% errors (2,3), which can lead to misalignments and ultimately impact the accuracy of downstream gene expression and alternative splicing analyses.

Methods for correcting long RNA sequencing reads fall into two classes: sequence-based and alignment-based. Early *sequence-based* correction methods were developed for long DNA reads used in genome assembly and aimed to correct the raw read sequences by aligning similar reads and correcting to their consensus (Canu, HGAP; (4,5)). For RNA reads, LoRMA (6) builds de Bruijn graphs of the sequences for varying k-mer sizes and uses the best path through the graph to correct sequences. IsOncorrect (7) clusters reads by sequence similarity and uses the consensus to correct sequences. DeSALT (8) is a two-step spliced alignment method that performs a seed-and-extend alignment to find exon cores, then refines their boundaries, improving the accuracy of splice junctions. Among *alignment-based* methods, TranscriptClean (9) corrects alignment artifacts such as micro-indels and non-canonical splice sites in co-localized read alignments. NanoSplicer (10) analyzes the raw electrical signals (“squiggles”) by aligning them to the candidate splice junctions using dynamic time warping. Lastly, transcript reconstruction tools developed their own built-in error correction, such as the highly accurate assembled ‘super-reads’ in StringTie (11), splice graph-based exon correction in IsoQuant (12), and exon boundary correction against a reference gene annotation in BAMBU (13), FLAIR (14), and the annotation-guided versions of the previous tools. However, standalone correction methods have shown limited performance, and built-in methods are restricted to the task at hand. Consequently, errors in the alignment, particularly at splice junctions, remain a critical and pressing challenge.

We developed SpliSync, a general-purpose tool for correcting splice sites in spliced alignments of Oxford Nanopore (ONT) sequences generated with diverse library preparation and sequencing technologies. Its underlying principle is that splice site correction in the alignment is sufficient for most transcriptomic applications, and more efficient and accurate than sequence correction. SpliSync combines a genomic language model (GLM), HyenaDNA (15), and a 1D U-net segmentation head, integrating genome sequence and alignment embeddings. Salient features include:

- Achieves high splice site accuracy, especially precision, 83.3-96.3% for human samples and 74.8-88.4% for cow and mouse, at a small loss of sensitivity (1-6%), while drastically outperforming competing methods;
- Increases the accuracy of splice sites by 27-194%, and of alternatively spliced intron groups by 26-330%, albeit the benefit for transcriptome assembly is limited due to their built-in filters and correction mechanisms;
- Performance is consistent across ONT RNA chemistries, library preparations, and sequencing technologies, and in multiple species;
- Does not require a reference annotation and can identify novel sites, making it suitable for species without high-quality annotation and for conditions where novel variation is expected, such as cancer samples.

SpliSync is written in Python using PyTorch. It is available free of charge under the GNU GPL 3.0 license from https://github.com/splicebox/SpliSync.

## SYSTEM AND METHODS

### The GLM-based model

#### Model description

SpliSync takes as input a set of alignments and the reference genomic sequence in a 10 kb window, and outputs at each genomic position the probability that it harbors a splice site. The underlying deep learning model integrates sequence and alignment information, combining a decoder-only genomic language model, HyenaDNA, with a feedforward neural network containing the alignment information, and a 1D U-Net (16) segmentation head for splice site prediction (**Figure 1**). To process the sequence input, we fine-tuned a pretrained HyenaDNA model with two Hyena layers, a width of 256, and a sequence length of 160k. The single-nucleotide tokenizer comprises ‘A’, ‘C’, ‘G’, ‘T’, and ‘N’, where ‘N’ serves as a padding or unspecified token. For alignment processing, numerical features derived from alignment counts and spliced read counts at each position are first variance-stabilized with log1p, then passed through the feedforward neural network to map to a higher-dimensional embedding space. The alignment-based embeddings are then concatenated with sequence-based embeddings produced by the Hyena model. The combined representation is passed to a 1D U-Net segmentation head composed of two encoder and two decoder blocks.

**Figure 1.**
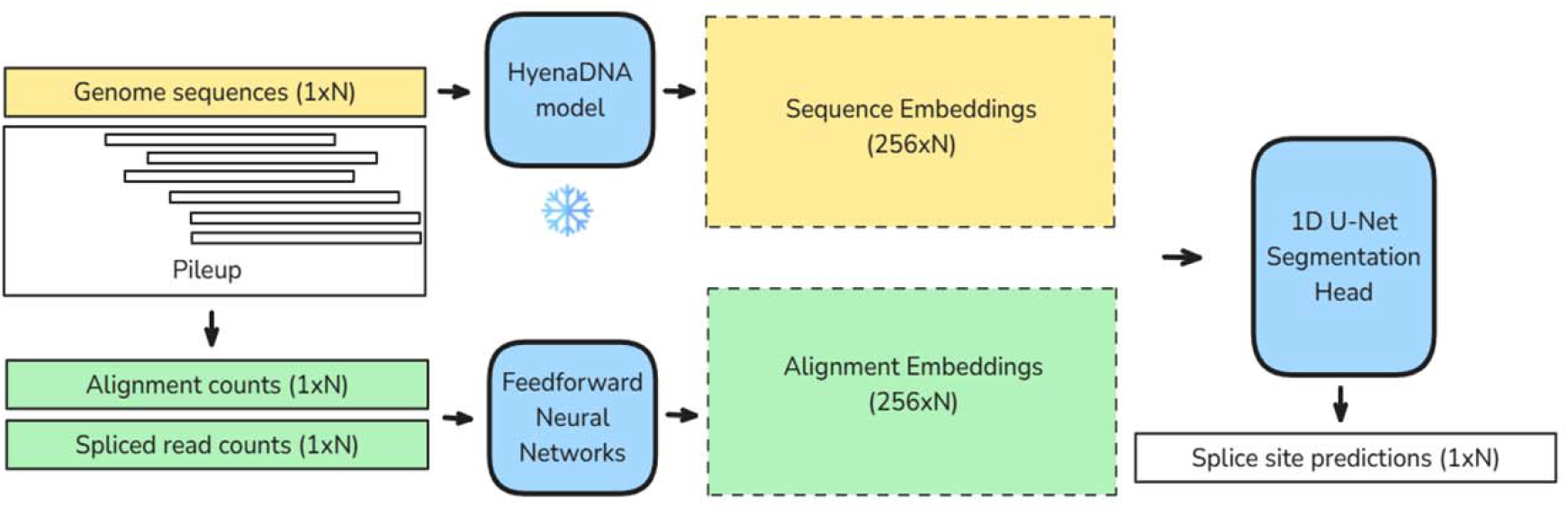
The SpliSync architecture. The SpliSync input consists of the genomic DNA sequence and alignment pile-up in that region. Sequence branch (top): the DNA sequence is processed by the pretrained and frozen HyenaDNA model to yield sequence embeddings. Alignment branch (bottom): from the pile-up, two coverage vectors are computed: one containing alignment counts and the other containing spliced-read counts. The coverage tracks are passed through feed-forward neural networks to produce alignment embeddings. The sequence and alignment embeddings are concatenated and input into a 1-D U-Net segmentation head, which outputs position-wise probabilities for splice-site prediction.

Overall, the channel dimensionality progresses from the initial concatenated input size, scaling up 2x and 4x, and subsequently scaling down to a final output projection of the unnormalized logits for binary splice site prediction at each nucleotide position. The SpliSync model has 210 million parameters.

#### Model training

We trained separate models for the R9.4 cDNA (*‘cdna’*), R9.4 direct RNA (‘*drna*’) and R10.4 cDNA (‘*r10*’) sequencing chemistries, as they exhibit differences in read coverage at the gene ends and in overall sequencing error rates (2). To train the ‘cdna’ model, we used NanoSim v.3.2.3 to simulate 3 million reads using the tool’s pretrained error model based on the expression profile of a human lymphoblastoid cell line, ERR14734450 (17). For the ‘drna’ model, we simulated 4 million reads using the error model and expression profile based on the human WTC-11 induced pluri-potent stem (iPS) cell line direct RNA data, ENCFF155CFF, from the Long-read RNA-Seq Genome Annotation Assessment Project (LRGASP) (18). Lastly, for the ‘r10’ model we simulated 4 million reads using the error model and expression profile of the human lymphoblastoid cell line, ERR14734450 (17). Details of the methods are presented in **Supplementary Method M1**.

Reads were mapped to the genome as described below. Regions with zero coverage or non-canonical splice sites were excluded. Collected data for each model was then split into disjoint subsets for training (chromosomes 1-19), validation (chromosome 22), and testing (chromosomes 21-22). The model was trained using the AdamW optimizer, with a learning rate 6e-4 and batch size of 64, for 5 epochs, using early stopping. To address the extreme class imbalance between the true splice sites and the rest of the genomic positions, we used a focal loss function with γ=2 and α=0.8, down-weighting the loss assigned to well-classified positions:

FL(p_t_) = − α (1 − p_t_)^γ^ log (p_t_), where γ ≥ 0 is the focusing parameter, α ∈ [0, 1] is a class-balancing weight, and p_t_ is the model’s estimated probability for the true class. Model training was conducted on a single compute node using four NVIDIA A100 GPUs, taking 12 hours to complete. More details are included in the **Supplementary Method M2**.

#### Post-processing

Splice-site predictions from SpliSync were used to correct raw read alignments by replacing native splice-junction coordinates with the closest model-predicted splice sites (within ±20 bp by default). Lastly, the sequence field in the alignment record was restored to the corresponding reference genome sequence.

### Benchmarking and evaluation

#### Sequence datasets

We applied the correction methods to six simulated and real datasets obtained as follows. Simulated human data was generated with NanoSim v.3.2.3 using the tool’s pretrained error model, corresponding to R9.4 cDNA sequencing with guppy basecalling, based on the expression profile of the lymphoblastoid cell line (LCL) CEPH1463 (NA12878) (3). Datasets of human, mouse and Chinese cattle ONT RNA sequences were obtained from GenBank: ERR14734531, human lymphocytes; ENCFF683TBO, mouse embryonic stem cell (mESCs); and CRR1887708, Chinese cattle, subcutaneous adipose tissue. In addition, the A549 cDNA replicate 6 (run 1) dataset from the Singapore Nanopore Expression Project (SGNEx) (19), derived from human lung carcinoma A549 cells, was obtained from the SGNEx consortium. Lastly, simulated mouse sequences conforming to the R10.4 technology were those reported in (12), obtained from https://zenodo.org/records/7611877. More details on the datasets are presented in **Table 1**.

**Table 1.**
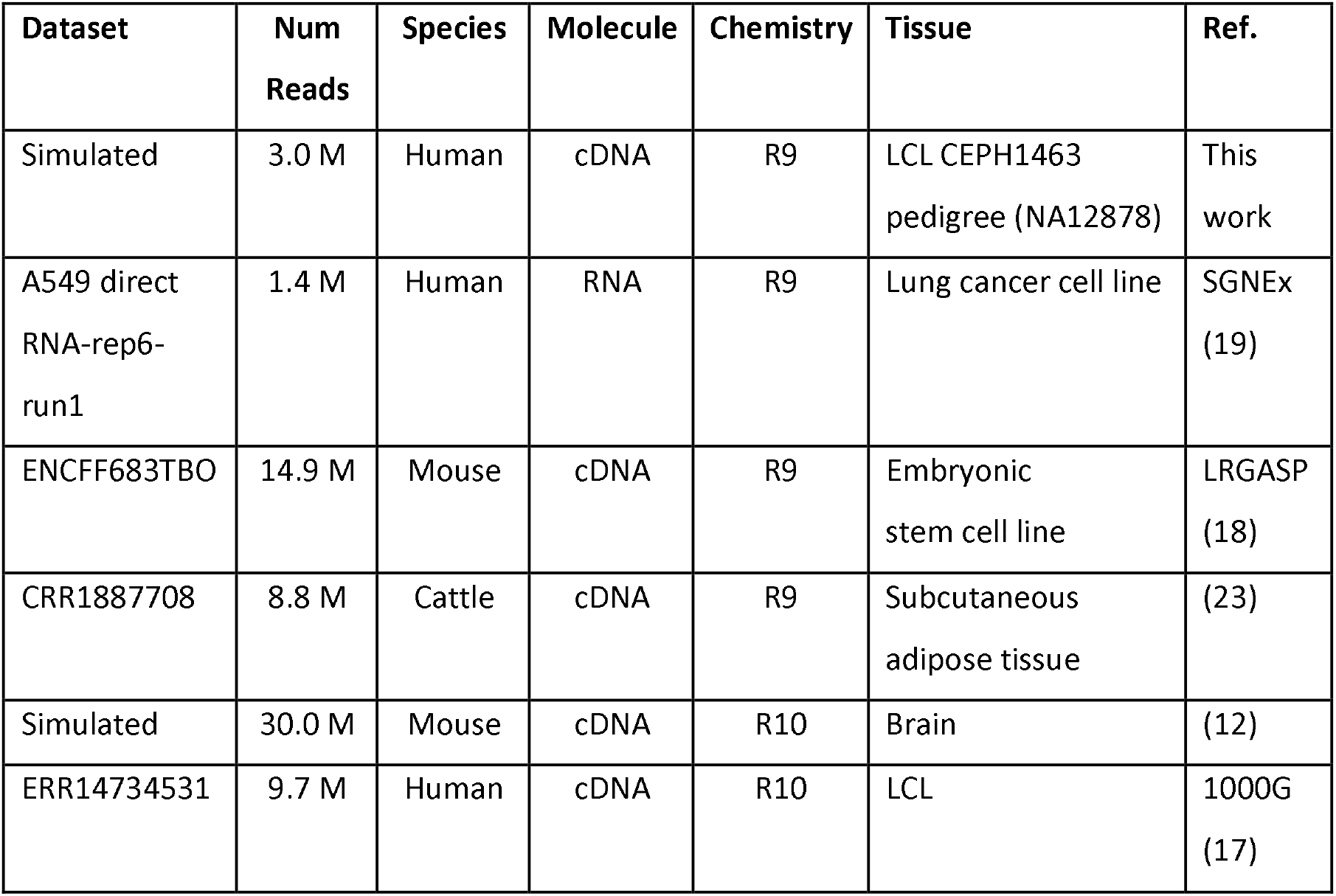
Selection of datasets used in the assessment.

#### Read processing and tools

Reads were mapped to the human GRCh38, mouse GRCm39, and *Bos taurus* ARS-UCD2.0 (GCA_002263795.4) genomes, respectively, using Minimap2 v.2.24 (20) with the options ‘-ax splice’ for ONT cDNA and ‘-ax splice -uf -k14’ for ONT direct RNA sequencing. For the simulated mouse data, which included a large fraction of polyA-tailed sequences, we removed the polyA tail using cutadapt v.5.2 with the options ‘-a A50’ and ‘—rc’, reverse complementing the output to restore the original orientation. Alignments were filtered to remove secondary and supplementary alignments with ‘samtools view’ using the flag filter ‘-F 3844’ before further processing. Splice junctions were extracted from the alignments using the utility ‘junc’ from the MntJULiP package (21,22). The error correction tools TranscriptClean v.2.1, LoRMA v.0.5, deSALT v1.5.6 and isONcorrect v0.0.8 were obtained from their GitHub or Sourceforge directory and applied, along with SpliSync, to all alignment datasets. LoRMA was slow, taking months to complete, while producing significantly suboptimal results, therefore was excluded from evaluations on datasets other than the simulated human data.

#### Applications

For the alternative splicing detection application, we used MntJULiP v.1.5.2 to extract introns from the spliced alignments and cluster them into ‘groups’ sharing a splice site (**Supplementary Figure S1**). Further, we filtered out introns supported by a single read in groups with 20 or more reads. For the transcriptome assembly application, assemblers FLAIR v.1.7.0, StringTie3 v.3.0.1, Bambu v3.4.0 and IsoQuant v3.10.0 were used to reconstruct transcripts from alignments, both *de novo* (reference-free) and with a reference gene annotation. For the simulated human dataset, to accurately assess the tools on a realistic scenario, we constructed the reference annotation by selecting one transcript not among the simulated isoforms, randomly, from each expressed gene. For all other datasets, we provided the tools with the species’ GENCODE annotations as reference. In the end, SQANTI3 v.5.2.1 was used to assess the accuracy of transcript reconstructions.

#### Evaluation scheme

Results from analyses on simulated data were evaluated against the set of simulated transcripts for both the human and mouse data, at the splice site and full-transcript levels. For analyses on real data, results were evaluated against GENCODE v.38 for the human datasets, GENCODE v.M26 for mouse, and ARS-UCD2.0 for cattle, using conventional measures of accuracy: sensitivity (recall), Sn = TP/(TP+FN); precision, Pr = TP/(TP+FP); and F1-value, F1 = 2*Sn*Pr/(Sn+Pr). For splice sites, an exact coordinate match was required. For alternative splicing groups, matched shared endpoints and at least two introns being present in the reference gene annotation were used to identify a match. For transcripts, categories were aggregated from the SQANTI3 output: *GeneVariant* = ‘FSM’ + ‘ISM’ + ‘NIC’ + ‘NNIC’, *Incompatible* = ‘Antisense’ + ‘Fusion’ + ‘Genic’ + ‘Intergenic’.

## MODEL AND EVALUATION

We first describe the model and calibration. Then, we benchmark SpliSync against existing read correction tools for the ability to correct splice sites. Lastly, we evaluate the impact of error correction on two downstream applications: alternative splicing detection and transcript reconstruction. To demonstrate the broad applicability and generalizability, we selected datasets produced with different ONT library preparation and sequencing technologies, and from different experiments, tissues, and species (**Table 1**).

### Overview of the model

We developed SpliSync, a deep learning method for accurate correction of splice sites in long-read RNA sequence alignments. SpliSync combines a genomic language model, HyenaDNA, with a 1D U-net segmentation head, integrating genome sequence and alignment embeddings. It takes as input the number of alignments and spliced alignments at each position in a 10 kb genomic interval, along with the nucleotide-level sequence in that interval. It outputs, at each position, a label indicating whether it represents a splice site: ‘positive’ (score ≥ 0.5) or ‘negative’ (score < 0.5). Additionally, we implemented SpliSync into a read corrector that outputs splice site-corrected alignments in BED or BAM format.

We trained this framework on simulated ONT data to create separate ‘images’ for the R9.4 cDNA (‘cdna’), R9.4 direct RNA (‘drna’), and R10.4 cDNA (‘r10’) technologies, as described in **Methods**. To determine what ‘image’ best suits a given library preparation and sequencing chemistry, we evaluated them on a collection of samples spanning different experiments (LRGASP, SGNEx), types of input material (cDNA, RNA), chemistries (R9.4, R10.4), sequencing technologies (R2C2, CapTrap, direct RNA, cDNA, direct cDNA), and species (human, fruitfly) (**Supplementary Figure S2**). (This collection is independent from that used for the program evaluation.) With a single exception, the fruitfly dataset, the R9.4 ‘*cdna*’ model achieved the best overall performance, albeit the results of all images were within a very close range (<1%). The correction improved the overall accuracy as measured by the F1-value for all sets, by 36% (human R9 cDNA LRGASP dataset) to 1% (human R10 cDNA LRGASP dataset), and precision by to 147% (human R9 cDNA LRGASP dataset) to 23% (human R9 R2C2 LRGASP, human R10 cDNA LRGASP datasets). Note that performance improvements may be obscured with the F1 metric due to artificially low sensitivity values, which are reflected from the entire universe of genes and transcripts, as represented in the reference annotation, and not only the expressed ones. The ‘cdna’ image consistently performed best and therefore was chosen as the program default, which was hereon applied to all analyses.

### Comparative evaluation of long RNA read error correction methods

We evaluated SpliSync and several error correction methods, including the sequence-based LoRMA (6) and isONcorrect (7), the spliced alignment corrector TranscriptClean (9), and the hierarchical splice junction-aware aligner deSALT (8), for the accuracy of splice site correction, using both simulated and real data. We used both simulated and real data spanning diverse technologies, including the R9.4 and R10.4 chemistries, cDNA and direct RNA sequencing, and various library construction protocols, sampled from multiple experiments, and from several species (human, mouse, Chinese cattle) (**Table 1**).

Performance was measured against the set of splice sites in the simulated transcripts for the simulated datasets, and against those extracted from the species-specific GENCODE gene annotations for the real data. While the latter does not provide an accurate measurement, since it overestimates the true positives by counting splice sites from paralogous genes and other alignment artifacts, it is a conventionally agreed-upon measure.

Across all datasets and categories, SpliSync was the only method to significantly improve prediction accuracy, with 5% to 71% increase in the F1-value over uncorrected data (**Figure 2**). Precision increased by 21% to 194%, reaching 83.3-96.3% on human, 81.8-88.4% on mouse, and 74.8% on cattle, with a very small loss in sensitivity of 1-6%. The improvement was seen across technologies. In particular, datasets obtained with the R10.4 chemistry, which produces higher-quality sequences with lower sequencing error rates, showed 64% and 46% increases in precision on the mouse-simulated and human CapTrap datasets, respectively.

**Figure 2.**
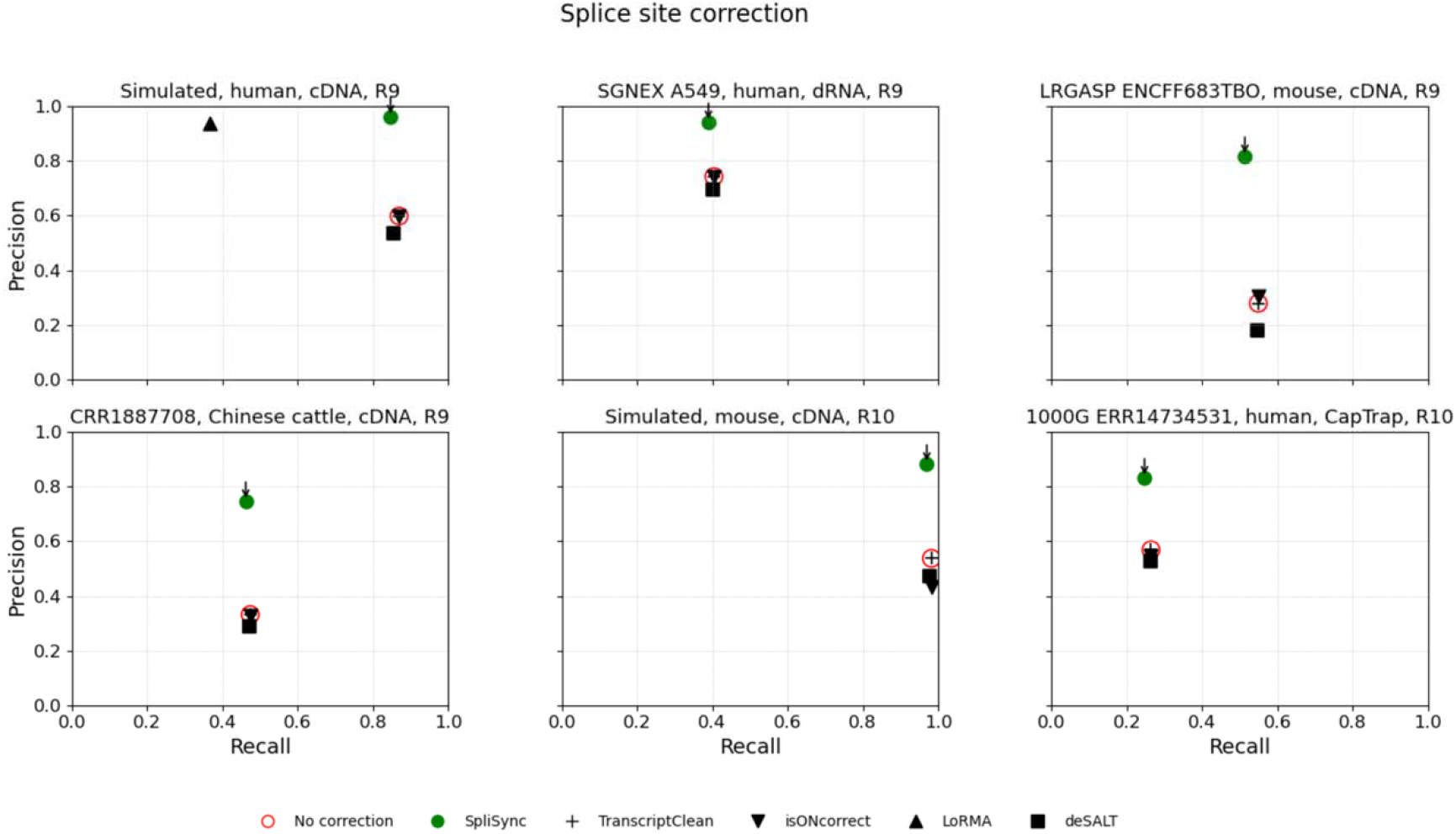
Performance evaluation of long RNA read error correction methods on splice site prediction. SpliSync was evaluated alongside competitor methods TranscriptClean, isONcorrect, LoRMA and deSALT on datasets produced with diverse ONT technologies, clockwise from the top left panel: simulated human data, cDNA, R9.4; SGNEx human A549 cell line data, dRNA, R9.4; LRGASP mouse data, cDNA, R9.4; Chinese cattle data, cDNA, R9.4; simulated mouse data, cDNA, R10.4; and 1000G human data, CapTrap, R10.4. Recall = TP/(TP+FN) is shown along the horizontal axis, and Precision = TP/(TP+FP) along the vertical one. A blank circle marks the uncorrected data, and arrows point to results produced by SpliSync.

### Application: Detection of alternative splicing variation

To assess the effects of splice site correction on alternative splicing detection, we constructed groups of competing introns that share an endpoint, thereby reflecting alternative splicing variations (**Supplementary Figure S1**). We then measured their accuracy before and after splice site correction on our data collection.

SpliSync was the best performer by a wide margin, while most programs failed to improve either the sensitivity or precision of alternative splicing variation detection for most of the datasets (**Figure 3**). SpliSync increased precision relative to uncorrected data by 13-196% (3-fold), depending on the dataset, with smaller changes in sensitivity, -17% to +6%, albeit the actual values vary among datasets. Notably, SpliSync improves both the sensitivity, by 6% and 9%, and the precision, by 13% and 330%, for the two R10 datasets. The largest improvement, by 192% and 330% (2.9-fold and 4.3-fold), is seen on the simulated data where the ground truth is known. In contrast, real data likely underestimates the precision improvement, as events classified as false positives may in fact represent novel alternative splicing variation and introns that are not present in the species-specific GENCODE database.

**Figure 3.**
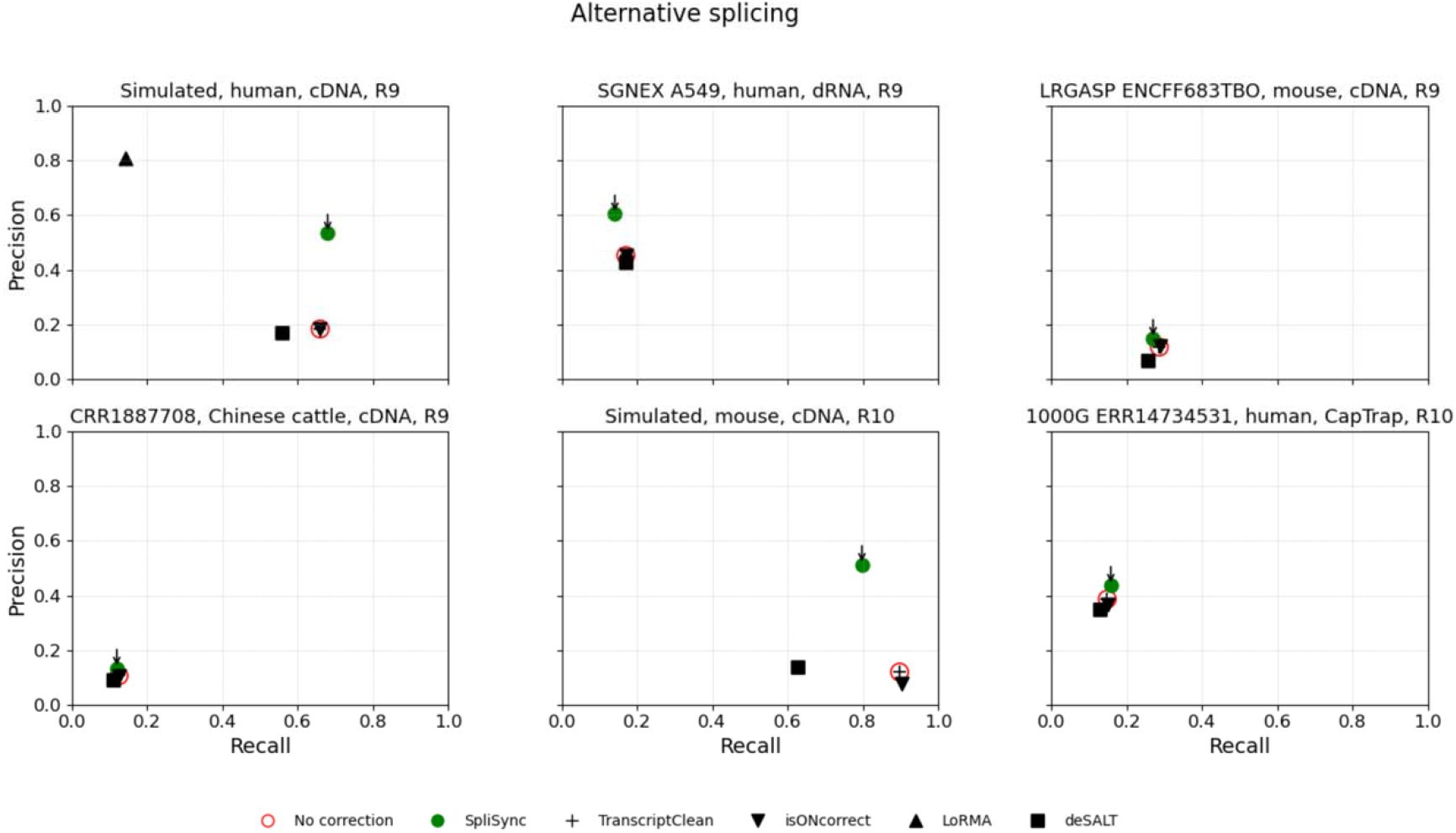
Impact evaluation of long RNA read error correction methods on alternative splicing detection. SpliSync was evaluated alongside competitor methods TranscriptClean, isONcorrect, LoRMA and deSALT on datasets produced with diverse ONT technologies, clockwise from the top left panel: simulated human data, cDNA, R9.4; SGNEx human A549 cell line data, dRNA, R9.4; LRGASP mouse data, cDNA, R9.4; Chinese cattle data, cDNA, R9.4; simulated mouse data, cDNA, R10.4; and 1000G human data, CapTrap, R10.4. Recall = TP/(TP+FN) is shown along the horizontal axis, and Precision = TP/(TP+FP) along the vertical one. A blank circle marks the uncorrected data, and an arrow points to results produced by SpliSync.

### Application: Transcriptome assembly

We assessed the effects of alignment correction on transcript assembly at the splice site and the transcript levels. We assembled the aligned reads, before and after correction with the aforementioned methods, using four popular long read transcript reconstruction methods, FLAIR, StringTie3, BAMBU, and IsoQuant, both with (‘| gtf’) and without (‘de novo’) using a reference gene annotation. For the simulated human data, to more realistically reflect the relationship between genes and transcripts in the reference annotation and the expressed transcriptome, we constructed a reference dataset by selecting one non-simulated transcript for each simulated gene. For all other datasets, assemblers were provided the species-specific GENCODE annotations. We measure the accuracy of predicted transcripts at the splice site and full-length transcript levels.

Surprisingly, error correction had little effect on performance (**Figure 4A** and **Supplementary Figures S3A-S7A**). Rather, splice site accuracy without and with alignment correction with the various methods clustered locally in the precision-recall (PRC) plot for each tool and application mode, namely, *de novo* or reference-assisted, denoted by ‘| gtf’. This clustering reflects the different design choices and internal algorithms developed by each assembler to filter and correct the error-prone input alignment data. Most methods were tuned for high precision (greater than 90%), except for FLAIR and BAMBU with reference annotations, and occasionally for StringTie3 and IsoQuant, whereas sensitivity varied widely even within the same dataset. In general, IsoQuant and BAMBU in *de novo* mode were the least sensitive, followed by IsoQuant| gtf, StringTie3, and FLAIR; FLAIR| gtf and StringTie3| gtf; and finally BAMBU| gtf.

**Figure 4.**
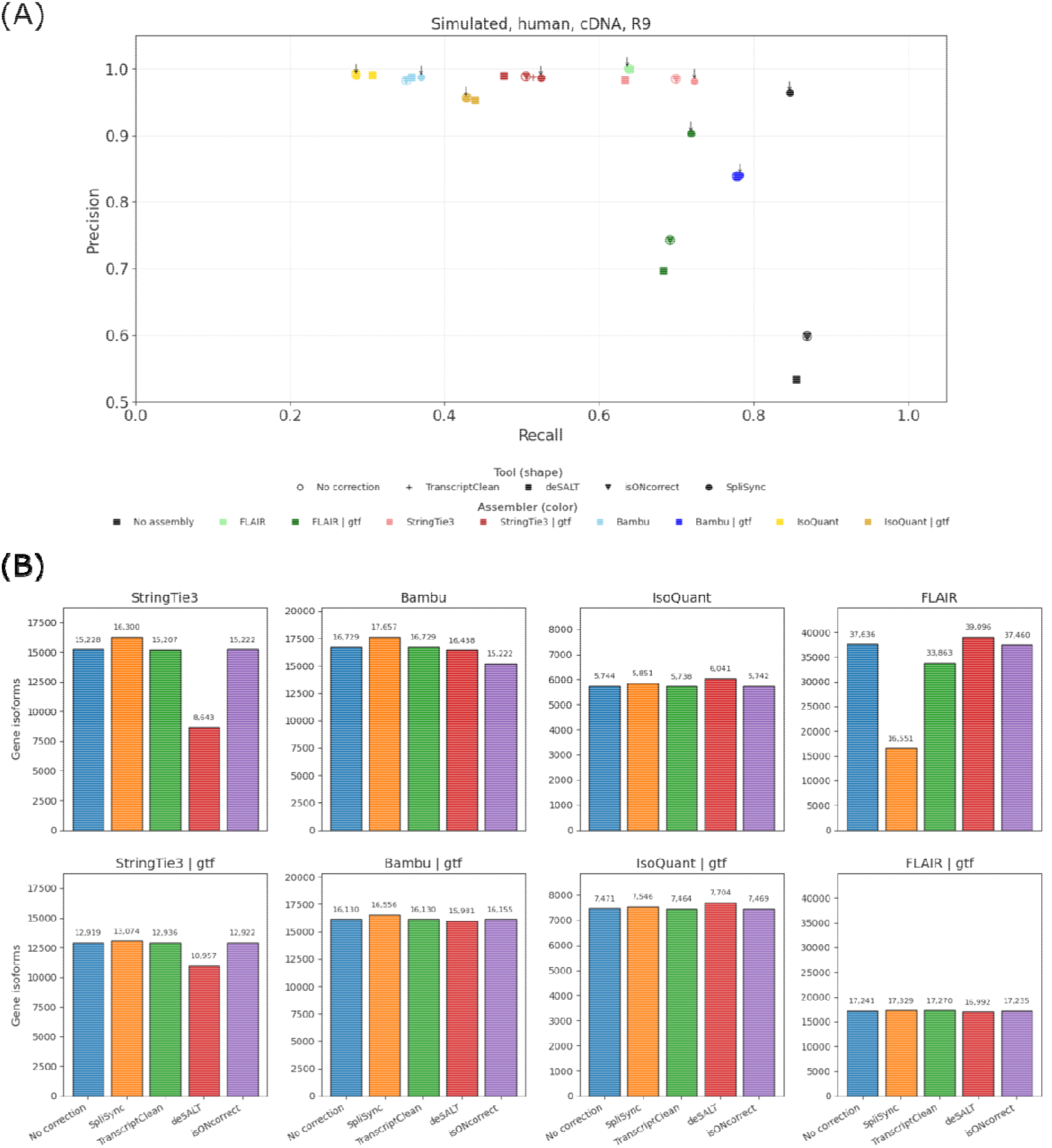
Impact evaluation of long RNA read error correction methods on transcript reconstruction, at the splice site (A) and transcript (B) levels, on the simulated human data. (A) Splice site-level evaluation. Assemblers (FLAIR, StringTie3, BAMBU and IsoQuant) are represented with colors, and correction methods (SpliSync, TranscriptClean, isONcorrect, LoRMA and deSALT) with different marks. Recall =TP/(TP+FN) is shown along the horizontal axis, and Precision = TP/(TP+FP)) along the vertical one. Unassembled data are shown in black, and arrows point to results produced by assemblers on SpliSync-corrected reads. (B) Transcript-level evaluation. Number of compatible transcripts ( *GeneVariant*) generated by each method combination, calculated from the SQANTI3 output (see Methods).

Annotation-assisted methods performed better than their *de novo* counterparts. However, as suggested by the simulated human data, this could be due to the tools’ over-reliance on reference annotations, leading to over-prediction. The most illustrative example is that of BAMBU| gtf, which ‘reconstructs’ almost the entire reference annotation and achieves close to perfect performance. In all cases, except for the artifactual outlier BAMBU| gtf, the splice site accuracy of the best performing methods remains significantly lower than that of the input splice sites corrected with SpliSync, suggesting the potential to develop new methods that capture additional transcripts and splice sites.

At the transcript level, splice site correction during pre-processing similarly resulted in only slight changes in the number of gene-compatible transcripts relative to the original alignments (**Figure 4B** and **Supplementary Figures S3B-S7B**). In the simulated human data, SpliSync led to modest increases (1-7%) for all assemblers except FLAIR and deSALT improved IsoQuant results by 3-5%, while all other correction tools led to a reduction in the number of compatible transcripts. Results were similar across all other datasets, with SpliSync showing small benefits for annotation-free assembly tasks and minimal effects when using a reference annotation, thereby supporting the earlier observations.

### Computational performance

For best performance, SpliSync should be run on GPU architectures; it is also CPU compatible. As an example, for the simulated human dataset comprising 3 million reads, preprocessing and model inference were completed in 13 and 15 min, respectively, using four NVIDIA A100-PCIE-40GB GPUs and a 128-core CPU, with a peak memory usage of 3.8GB RAM on a 2.1 GHz Intel Xeon Gold server. In contrast, CPU-only inference took 568 min, indicating an approximately 38-fold speed-up when GPUs are used.

## IMPLEMENTATION AND AVAILABILITY

SpliSync is implemented in Python using the PyTorch package and is freely available at https://github.com/splicebox/SpliSync. An archived version, including model weights, is available from Zenodo (DOI: 10.5281/zenodo.20148684).

## DISCUSSION AND CONCLUSIONS

While long-read sequencing offers the potential to resolve full-length isoforms, its effectiveness is often limited by high error rates that lead to misaligned splice junctions. We developed SpliSync, a genomic language model-driven method that significantly improves the precision of splice site annotations in long read spliced alignments. SpliSync integrates the HyenaDNA decoder-only architecture with a 1D U-Net segmentation head, leveraging both genomic language embeddings and local alignment pile-up evidence to refine splice-site predictions.

SpliSync significantly increases splice site accuracy by prioritizing precision, enabling more reliable identification of alternative splicing variation. SpliSync was validated across Oxford Nanopore R9 and R10 chemistries, with diverse library preparations, and across multiple species, demonstrating robust accuracy.

SpliSync can identify splice sites directly from the input alignments, without relying on a reference annotation, and can identify novel sites. Therefore, it is well-suited for application in species without a comprehensive annotation or in conditions expected to exhibit novel splice variations, such as cancer.

SpliSync substantially improved the accuracy of splice sites in ONT long RNA read alignments, in particular precision, by 27-194%. While it also significantly improved accuracy in detecting alternative splicing in the form of intron groups, precision remains below the desirable level, at <60%. Based on observation, most false positive events occur at known introns created by artifactual variations around the non-shared intron endpoint, which remains a systematic error. Further model training coupled with heuristics on read support for introns could help resolve these. Additionally, while SpliSync’s current model is suitable for mammalian species and potentially across the broader vertebrate subphylum, application to more divergent species and plants will require alternative genomic language models and training on specific lineages. Finally, some application-specific tools, such as transcript assemblers, have developed their own internal correction mechanisms and saw little benefit from the correction. However, the significant gap between the SpliSync-corrected input and the tools’ generated splice sites suggests that new and more sensitive assembly methods could be developed to leverage the highly accurate input.

SpliSync produces a list of splice sites and a set of corrected alignments. It can be used standalone as a general-purpose preprocessing tool ahead of other applications, including clustering, alternative splicing discovery, quantification, and differential expression and splicing analysis, or can be integrated directly into accurate and efficient transcriptomic analysis tools.

## DATA AVAILABILITY

Simulated human long RNA sequencing reads are available from Zenodo (DOI: 10.5281/zenodo.20148684). All other sequencing datasets are available from the SRA and as referenced in the text. Analysis scripts can be obtained from the SpliSync’s GitHub repository, and results of the analyses are available on Zenodo (DOI: 10.5281/zenodo.20148684).

## ACKNOWLEDGMENTS

Computations were performed on the Advanced Research Computing at Hopkins (ARCH) facility supported by the National Science Foundation [OAC 1920103]. This manuscript is the result of funding in whole or in part by the National Institutes of Health (NIH). It is subject to the NIH Public Access Policy. Through acceptance of this federal funding, NIH has been given a right to make this manuscript publicly available in PubMed Central upon the Official Date of Publication, as defined by NIH.

## REFERENCES

1. Ament, I.H., DeBruyne, N., Wang, F. and Lin, L. (2025) Long-read RNA sequencing: A transformative technology for exploring transcriptome complexity in human diseases. Mol Ther, 33, 883–894.

2. Liu-Wei, W., van der Toorn, W., Bohn, P., Holzer, M., Smyth, R.P. and von Kleist, M. (2024) Sequencing accuracy and systematic errors of nanopore direct RNA sequencing. BMC Genomics, 25, 528.

3. Workman, R.E., Tang, A.D., Tang, P.S., Jain, M., Tyson, J.R., Razaghi, R., Zuzarte, P.C., Gilpatrick, T., Payne, A., Quick, J. et al. (2019) Nanopore native RNA sequencing of a human poly(A) transcriptome. Nat Methods, 16, 1297–1305.

4. Koren, S., Walenz, B.P., Berlin, K., Miller, J.R., Bergman, N.H. and Phillippy, A.M. (2017) Canu: scalable and accurate long-read assembly via adaptive k-mer weighting and repeat separation. Genome Res, 27, 722–736.

5. Chin, C.S., Alexander, D.H., Marks, P., Klammer, A.A., Drake, J., Heiner, C., Clum, A., Copeland, A., Huddleston, J., Eichler, E.E. et al. (2013) Nonhybrid, finished microbial genome assemblies from long-read SMRT sequencing data. Nat Methods, 10, 563–569.

6. Salmela, L., Walve, R., Rivals, E. and Ukkonen, E. (2017) Accurate self-correction of errors in long reads using de Bruijn graphs. Bioinformatics, 33, 799–806.

7. Sahlin, K. and Medvedev, P. (2021) Error correction enables use of Oxford Nanopore technology for reference-free transcriptome analysis. Nat Commun, 12, 2.

8. Liu, B., Liu, Y., Li, J., Guo, H., Zang, T. and Wang, Y. (2019) deSALT: fast and accurate long transcriptomic read alignment with de Bruijn graph-based index. Genome Biol, 20, 274.

9. Wyman, D. and Mortazavi, A. (2019) TranscriptClean: variant-aware correction of indels, mismatches and splice junctions in long-read transcripts. Bioinformatics, 35, 340–342.

10. You, Y., Clark, M.B. and Shim, H. (2022) NanoSplicer: accurate identification of splice junctions using Oxford Nanopore sequencing. Bioinformatics, 38, 3741–3748.

11. Shumate, A., Wong, B., Pertea, G. and Pertea, M. (2022) Improved transcriptome assembly using a hybrid of long and short reads with StringTie. PLoS Comput Biol, 18, e1009730.

12. Prjibelski, A.D., Mikheenko, A., Joglekar, A., Smetanin, A., Jarroux, J., Lapidus, A.L. and Tilgner, H.U. (2023) Accurate isoform discovery with IsoQuant using long reads. Nat Biotechnol, 41, 915–918.

13. Chen, Y., Sim, A., Wan, Y.K., Yeo, K., Lee, J.J.X., Ling, M.H., Love, M.I. and Goke, J. (2023) Context-aware transcript quantification from long-read RNA-seq data with Bambu. Nat Methods, 20, 1187–1195.

14. Tang, A.D., Soulette, C.M., van Baren, M.J., Hart, K., Hrabeta-Robinson, E., Wu, C.J. and Brooks, A.N. (2020) Full-length transcript characterization of SF3B1 mutation in chronic lymphocytic leukemia reveals downregulation of retained introns. Nat Commun, 11, 1438.

15. Nguyen, E., Poli, M., Faizi, M., Thomas, A., Birch-Sykes, C., Wornow, M., Patel, A., Rabideau, C., Massaroli, S., Bengio, Y. et al. (2023) HyenaDNA: Long-range genomic sequence modeling at single nucleotide resolution. NeurIPS2023. ArXiv:2306.15794.

16. Ronneberger, O., Fischer, P. and Brox, T. (2015). U-Net: Convolutional Networks for Biomedical Image Segmentation. Medical Image Computing and Computer-Assisted Intervention – MICCAI 2015, Springer International Publishing, Cham, pp. 234–241.

17. Clavell-Revelles, P., Reese, F., Carbonell-Sala, S., Degalez, F., Arnan, C., Oliveros, W., Palumbo, E., Perteghella, T., Guigo, R. and Mele, M. (2025) Long-read transcriptomics of a diverse human cohort reveals ancestry bias in gene annotation. Nat Commun, 16, 10194.

18. Pardo-Palacios, F.J., Wang, D., Reese, F., Diekhans, M., Carbonell-Sala, S., Williams, B., Loveland, J.E., De Maria, M., Adams, M.S., Balderrama-Gutierrez, G. et al. (2024) Systematic assessment of long-read RNA-seq methods for transcript identification and quantification. Nat Methods, 21, 1349–1363.

19. Chen, Y., Davidson, N.M., Wan, Y.K., Yao, F., Su, Y., Gamaarachchi, H., Sim, A., Patel, H., Low, H.M., Hendra, C. et al. (2025) A systematic benchmark of Nanopore long-read RNA sequencing for transcript-level analysis in human cell lines. Nat Methods, 22, 801–812.

20. Li, H. (2018) Minimap2: pairwise alignment for nucleotide sequences. Bioinformatics, 34, 3094–3100.

21. Lui, W.W., Yang, G., He, Z. and Florea, L. (2025) MntJULiP and Jutils: differential splicing analysis of RNA-seq data with covariates. NAR Genom Bioinform, 7, lqaf140.

22. Yang, G., Sabunciyan, S. and Florea, L. (2022) Comprehensive and scalable quantification of splicing differences with MntJULiP. Genome Biol, 23, 195.

23. Fang, W., Yang, S., Li, X., Nanaei, A., Liu, Y., Cao, Y., Xiao, C., Li, X., Jin, H., Zhao, Y. et al. (2025) Long-read sequencing uncovers key regulatory genes involved in the differentiation of preadipocytes of Chinese red steppe cattle. Sci Rep, 15, 29459.

