## Supplementary Figures and Tables for "SpliSync: Genomic language model-driven splice site correction of long RNA sequencing reads"

SUPPLEMENTARY MATERIAL FOR THE ARTICLE:
"SPLISYNC: GENOMIC LANGUAGE MODEL-DRIVEN SPLICE SITE CORRECTION OF LONG RNA SEQUENCING READS"

BY W.W. LUI and L. FLOREA

**TABLE OF CONTENTS:**

**Supplementary Methods:**

**Method M1.** Training dataset construction and pre-processing.
**Method M2.** Data simulation for SpliSync 'image' generation.

**Supplementary Figures:**

**Figure S1.** Representation of intron-based alternative splicing 'groups'.
**Figure S2.** Comparison and calibration of SpliSync's 'cdna', 'drna, and 'r10' image models.
**Figure S3.** Impact evaluation of long RNA read error correction methods on transcript reconstruction on the SGNEx human A549 cell line, dRNA, R9.4 dataset.
**Figure S4.** Impact evaluation of long RNA read error correction methods on transcript reconstruction on the LRGASP mouse, cDNA, R9.4 dataset.
**Figure S5.** Impact evaluation of long RNA read error correction methods on transcript reconstruction on the Chinese cattle, cDNA, R9.4 dataset.
**Figure S6.** Impact evaluation of long RNA read error correction methods on transcript reconstruction on the simulated mouse data, cDNA, R10.4 dataset.
**Figure S7.** Impact evaluation of long RNA read error correction methods on transcript reconstruction on the 1000G human LCL, CapTrap, R10.4 dataset.

### SUPPLEMENTARY METHODS

#### Supplementary Method M1. Training dataset construction and pre-processing

To train and evaluate the splice-site prediction model, we developed an automated data curation pipeline to extract sequence and alignment features from reference FASTA and corresponding BAM files. Alongside the primary nucleotide sequences, we generated quantitative alignment profiles to provide the model with empirical read coverage context. For each genomic position within the target regions, we used pysam v.0.22.1 to parse alignment pileups and extract two core features: total read depth (the count of aligned reads covering the position, excluding deletions) and spliced read count (the count of reads skipping the reference base, indicating a splice junction). These features were aggregated into a two-dimensional matrix for each target region. The extracted sequences and corresponding pileup matrices were systematically segmented into uniform chunks of N base pairs (default N = 10,000). Sequences shorter than this window size, including terminal chunks, were padded with 'N' nucleotides. The extraction pipeline was parallelized using the Dask v.2023.5.0 distributed computing framework. Following model inference, predictions were further refined by applying zero-read support filters and merging adjacent positive calls within a default 20-bp window. These inputs form the foundation for downstream training, validation and benchmarking analyses presented in this study.

#### Supplementary Method M2. Data simulation for SpliSync 'image' generation

To train NanoSim's sequencing error models (cDNA R9.4, direct RNA R9.4, cDNA R10.4), we randomly subsampled 1M reads from the corresponding real dataset and ran 'read_analysis.py transcriptome' with the option '--no_intron_retention'. Raw FASTQ reads from the target expression dataset were first aligned to GENCODE v.38 transcripts with minimap2 v.2.24 (option'-ax map-ont'). Transcript abundances were then estimated with Salmon v.1.9.0 in alignment-based mode (option '--ont'). Only transcripts annotated as "protein_coding" or "lncRNA" were retained for downstream simulation. For simulating both cDNA and direct RNA (dRNA) libraries with NanoSim's simulator.py, the basecalling error profile was modelled after Guppy (--guppy). For the dRNA dataset, we specifically enabled uracil modeling (--uracil). These configurations were used to generate the simulated sequence data for training and evaluating SpliSync's 'images' (*'cdna', 'drna', 'r10'*).

### SUPPLEMENTARY FIGURES

**Supplementary Figure S1.** Representation of intron-based alternative splicing 'groups'. A 'group' is formed of all introns obtained from the input spliced alignments and that share an endpoint.

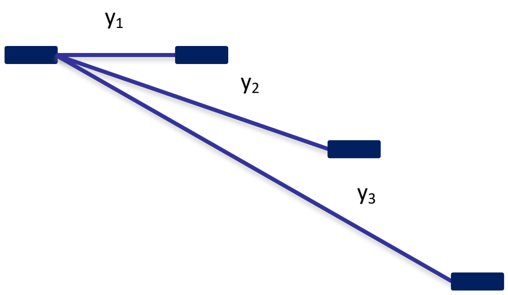

**Supplementary Figure S2.** Comparison and calibration of SpliSync's 'cdna', 'drna, and 'r10' image models. We evaluated the performance of the three SpliSync images on an independent set of samples: LRGASP ENCFF263YFG (human, R9, cDNA), ENCFF771DIX (human, R9, direct RNA), ENCFF377IEH (human, R9, R2C2), ENCFF934KDM (human, R9, CapTrap), and SRR23881262 (human, R10, cDNA); SGNEx A549 (human, R9, cDNA) and A549 (human, R9, direct cDNA); and GenBank SRA SRR35042122 (fruitfly R9, cDNA). While the three models had close performance for each of the datasets, the 'cdna' image trained on R9.4 simulated data had the best overall performance as measured by F1-val= 2*Sn*Pr/(Sn+Pr), for each dataset except for the fruitfly sample, on which it ranked as a very close second.

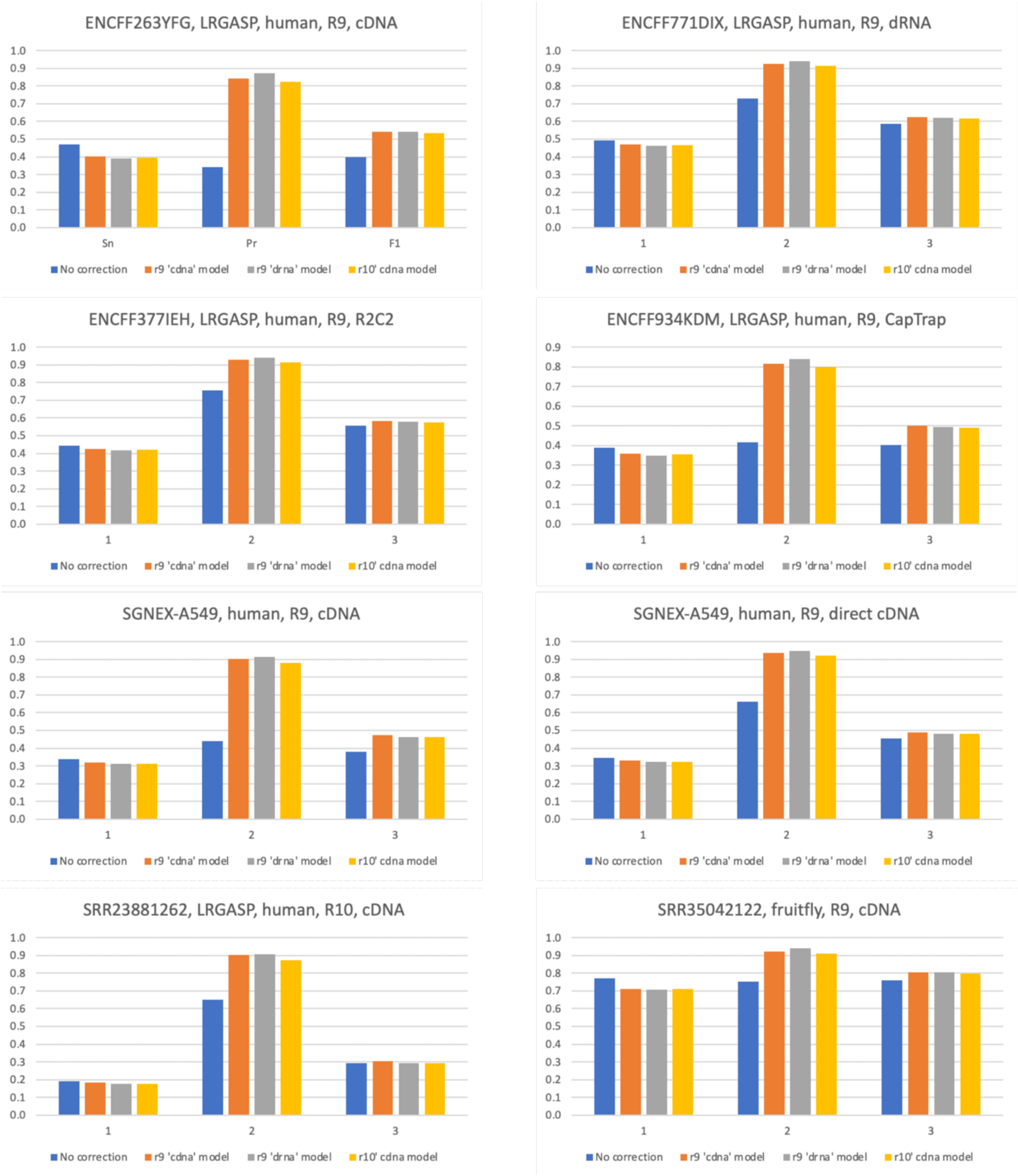

**Supplementary Figure S3.** Impact evaluation of long RNA read error correction methods on transcript reconstruction on the SGNEx human A549 cell line, dRNA, R9.4 dataset. (A) Splice site-level evaluation: Assemblers (FLAIR, StringTie3, BAMBU and IsoQuant) are represented with colors, and correction methods (SpliSync, TranscriptClean, isONcorrect, LoRMA and deSALT) with different marks. Recall =TP/(TP+FN) is shown along the horizontal axis, and Precision = TP/(TP+FP)) along the vertical one. Unassembled data are shown in black, and arrows point to results produced by assemblers on SpliSync-corrected reads. (B) Transcript-level evaluation: Number of compatible transcripts (*GeneVariant*) generated by each method combination, calculated from the SQANTI3 output.

(A)

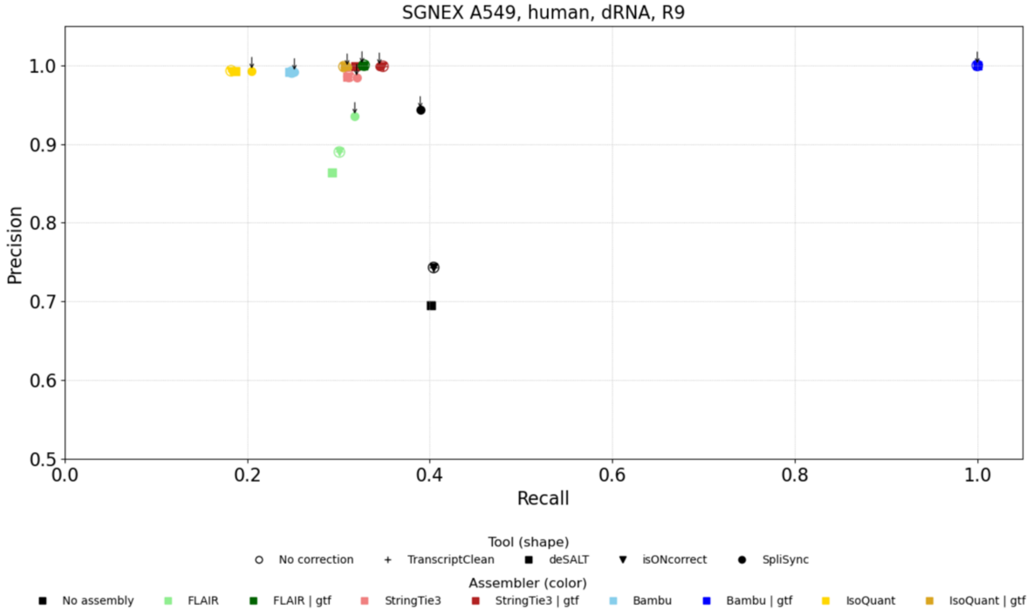


(B)

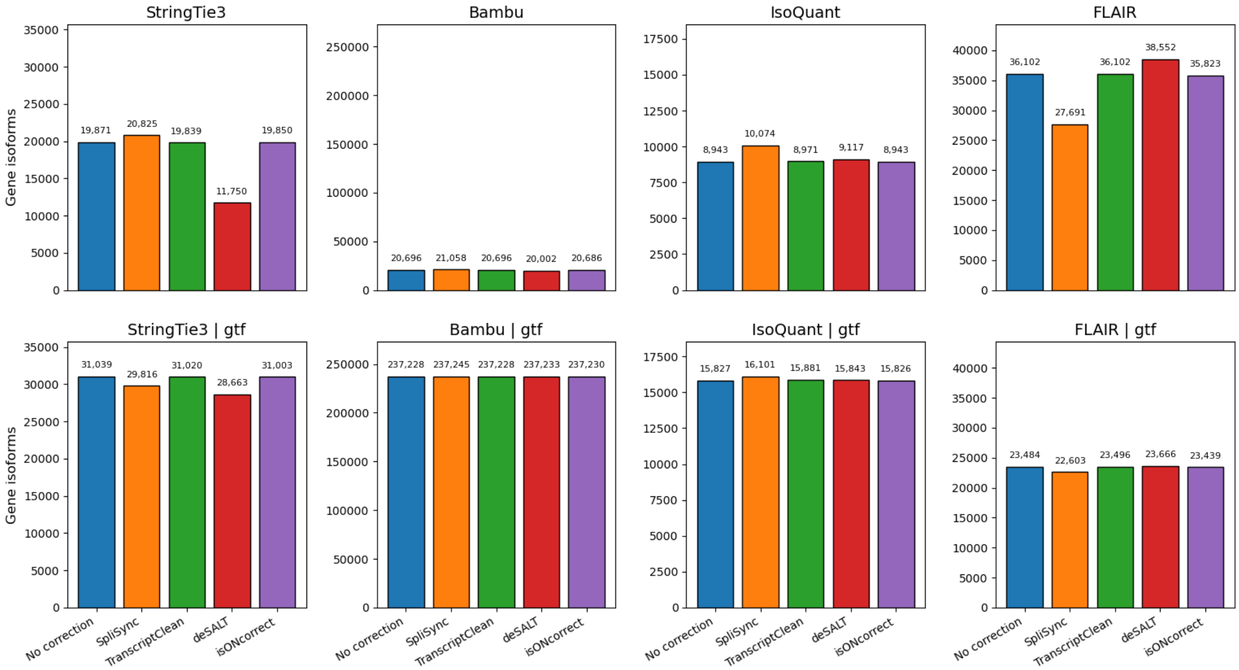

**Supplementary Figure S4.** Impact evaluation of long RNA read error correction methods on transcript reconstruction on the LRGASP mouse, cDNA, R9.4 dataset. (A) Splice site-level evaluation, and (B) Assemblers (FLAIR, StringTie3, BAMBU and IsoQuant) are represented with colors, and correction methods (SpliSync, TranscriptClean, isONcorrect, LoRMA and deSALT) with different marks. Recall =TP/(TP+FN) is shown along the horizontal axis, and Precision = TP/(TP+FP)) along the vertical one. Unassembled data are shown in black, and arrows point to results produced by assemblers on SpliSync-corrected reads. (B) Transcript-level evaluation: Number of compatible transcripts (*GeneVariant*) generated by each method combination, calculated from the SQANTI3 output.

(A)

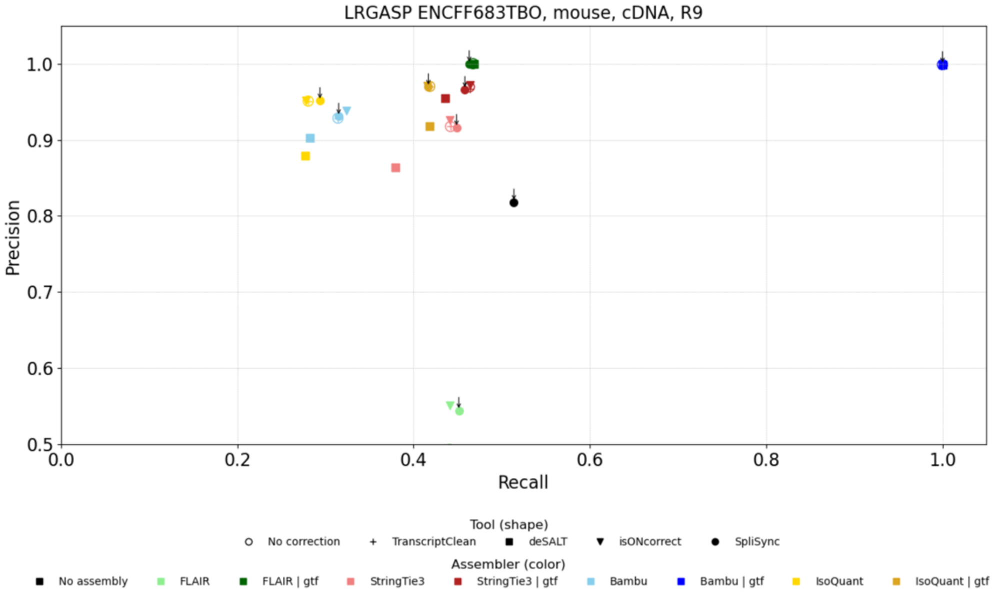


(B)

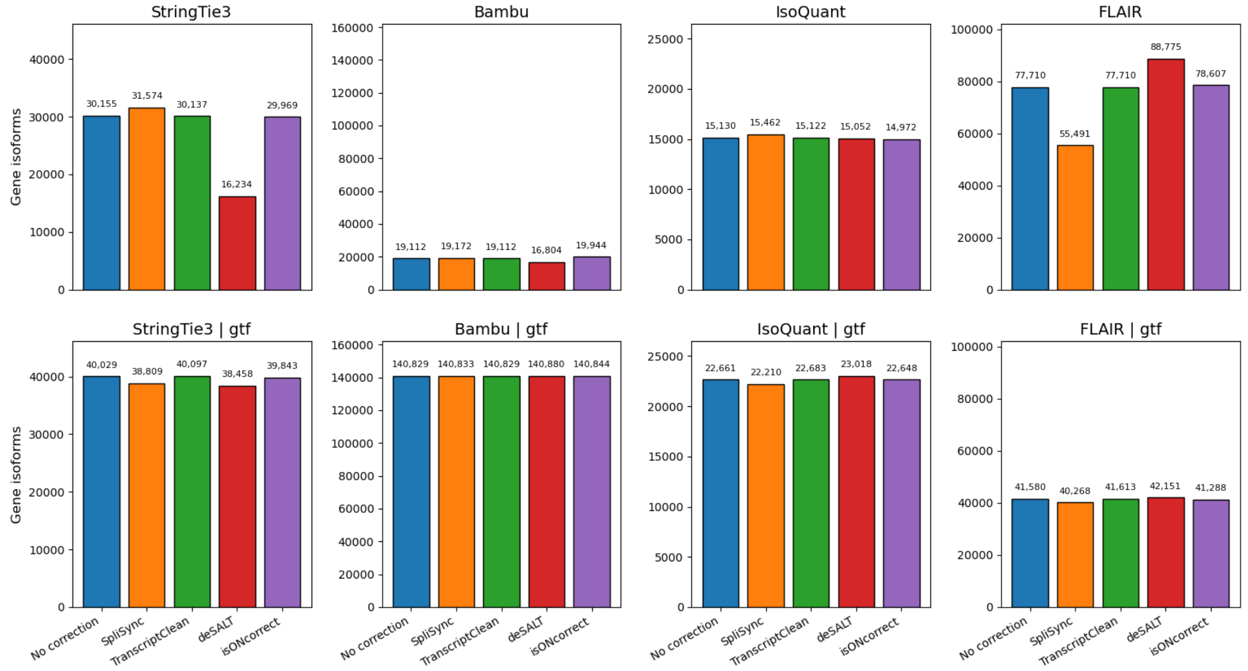

**Supplementary Figure S5.** Impact evaluation of long RNA read error correction methods on transcript reconstruction on the Chinese cattle, cDNA, R9.4 dataset. A) Splice site-level evaluation, and (B) Assemblers (FLAIR, StringTie3, BAMBU and IsoQuant) are represented with colors, and correction methods (SpliSync, TranscriptClean, isONcorrect, LoRMA and deSALT) with different marks. Recall =TP/(TP+FN) is shown along the horizontal axis, and Precision = TP/(TP+FP)) along the vertical one. Unassembled data are shown in black, and arrows point to results produced by assemblers on SpliSync-corrected reads. (B) Transcript-level evaluation: Number of compatible transcripts (*GeneVariant*) generated by each method combination, calculated from the SQANTI3 output.

(A)

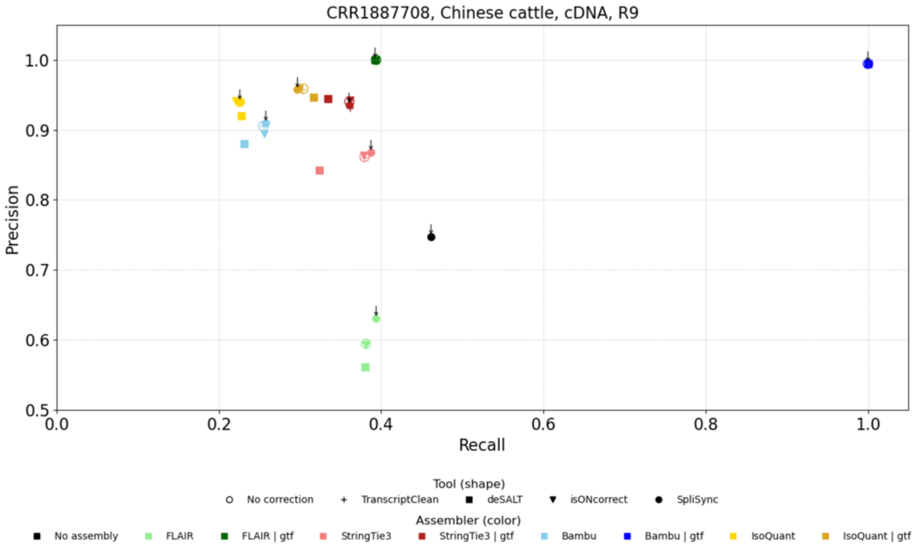


(B)

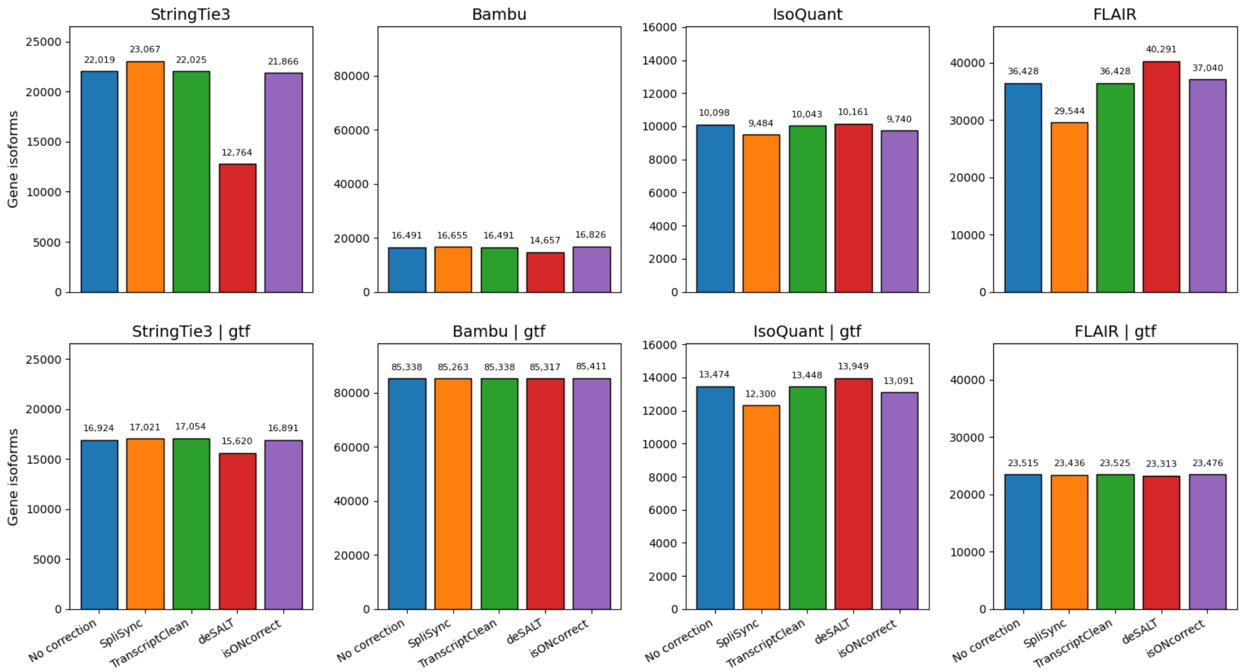

**Supplementary Figure S6.** Impact evaluation of long RNA read error correction methods on transcript reconstruction on the simulated mouse data, cDNA, R10.4 dataset. (A) Splice site-level evaluation, and (B) Assemblers (FLAIR, StringTie3, BAMBU and IsoQuant) are represented with colors, and correction methods (SpliSync, TranscriptClean, isONcorrect, LoRMA and deSALT) with different marks. Recall =TP/(TP+FN) is shown along the horizontal axis, and Precision = TP/(TP+FP)) along the vertical one. Unassembled data are shown in black, and arrows point to results produced by assemblers on SpliSync-corrected reads. (B) Transcript-level evaluation: Number of compatible transcripts (*GeneVariant*) generated by each method combination, calculated from the SQANTI3 output.

(A)

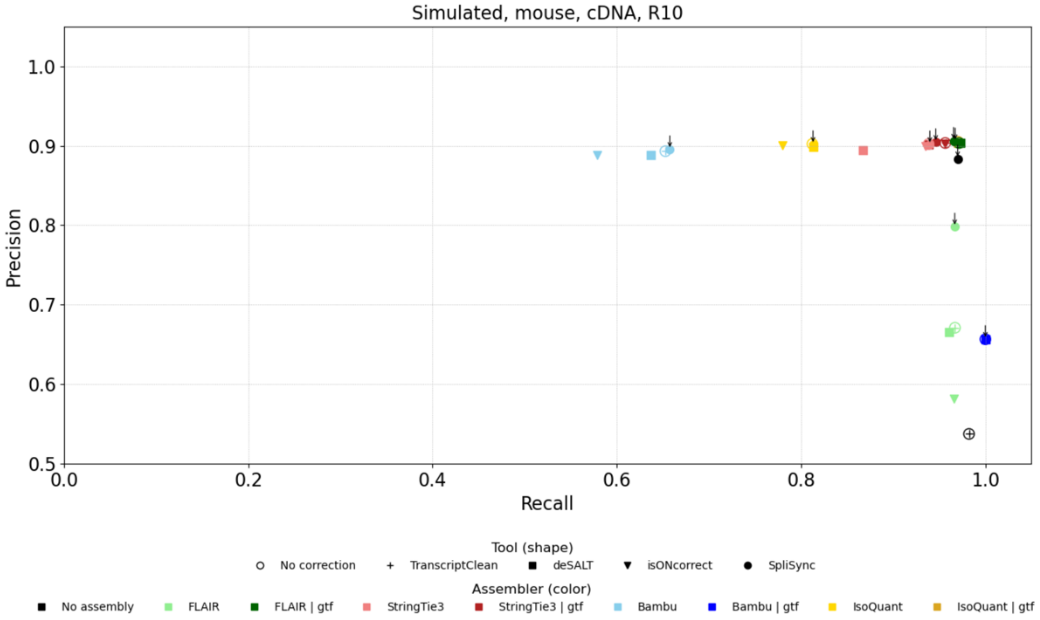


(B)

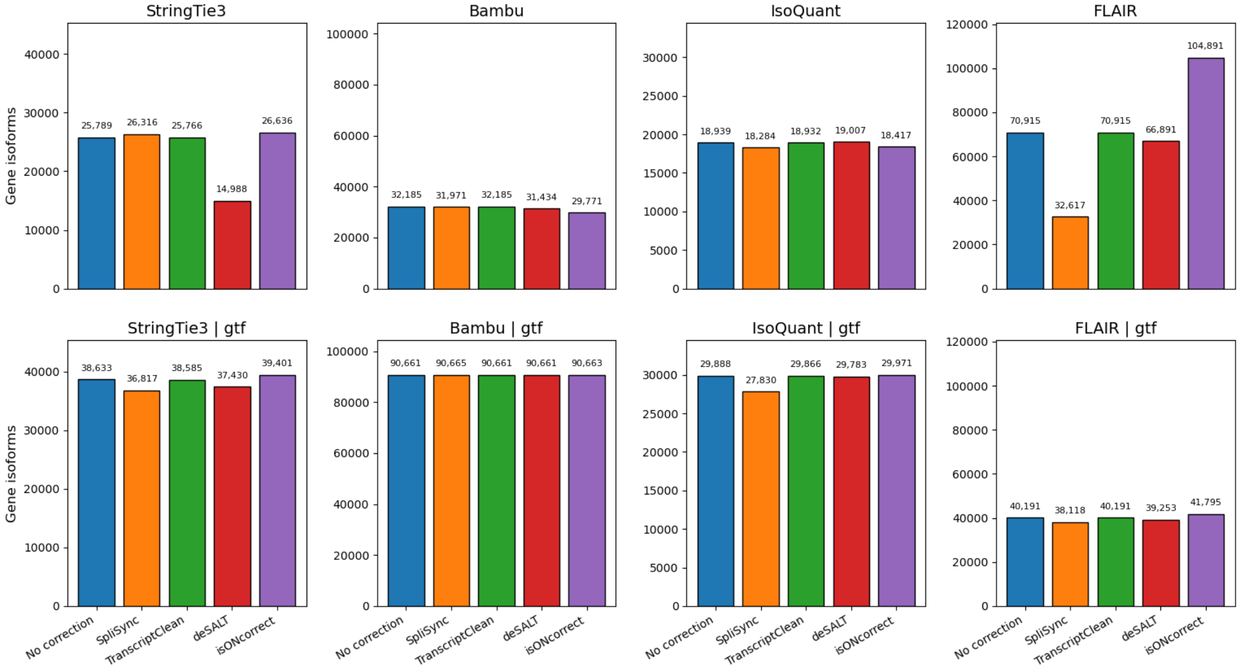

**Supplementary Figure S7.** Impact evaluation of long RNA read error correction methods on transcript reconstruction on the 1000G human LCL, CapTrap, R10.4 dataset. (A) Splice site-level evaluation, and (B) Assemblers (FLAIR, StringTie3, BAMBU and IsoQuant) are represented with colors, and correction methods (SpliSync, TranscriptClean, isONcorrect, LoRMA and deSALT) with different marks. Recall =TP/(TP+FN) is shown along the horizontal axis, and Precision = TP/(TP+FP)) along the vertical one. Unassembled data are shown in black, and arrows point to results produced by assemblers on SpliSync-corrected reads. (B) Transcript-level evaluation: Number of compatible transcripts (*GeneVariant*) generated by each method combination, calculated from the SQANTI3 output.

(A)

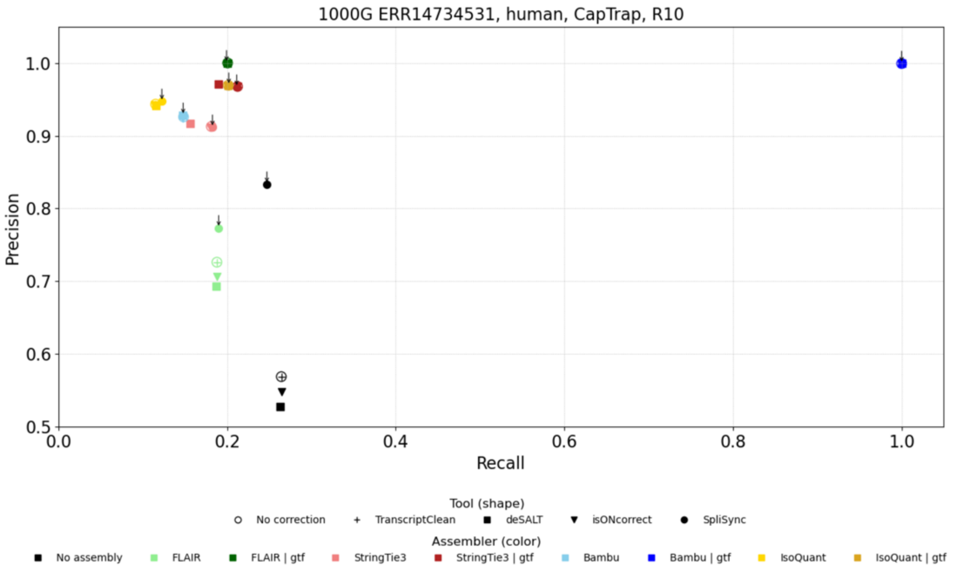


(B)

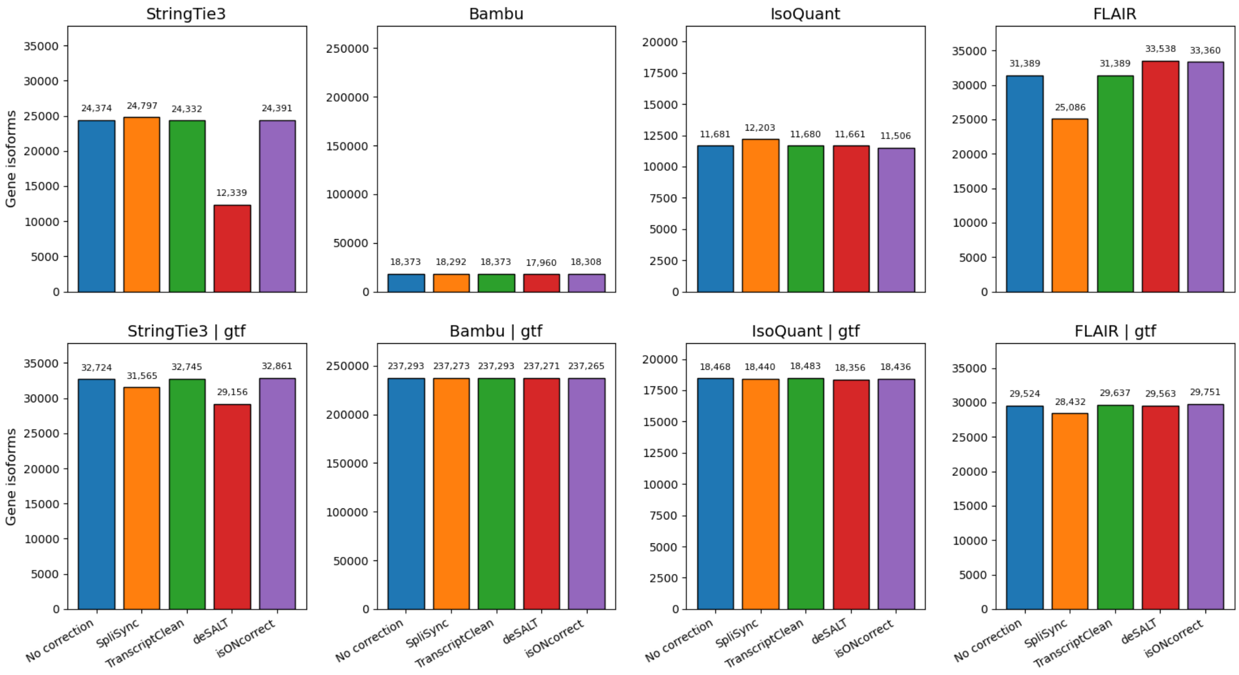
